# Nucleation and stability of branched versus linear Arp2/3-generated actin filaments

**DOI:** 10.1101/2022.05.06.490861

**Authors:** Luyan Cao, Foad Ghasemi, Michael Way, Antoine Jégou, Guillaume Romet-Lemonne

## Abstract

Activation of the Arp2/3 complex by VCA-domain-bearing NPFs results in the formation of ‘daughter’ actin filaments branching off the sides of pre-existing ‘mother filaments.’ Alternatively, when stimulated by SPIN90, Arp2/3 directly nucleates ‘linear’ actin filaments. Uncovering the similarities and differences of these two activation mechanisms is fundamental to understanding the regulation and function of Arp2/3. Analysis of individual filaments reveals that the catalytic VCA domain of WASP, N-WASP and WASH, accelerate the Arp2/3-mediated nucleation of linear filaments by SPIN90, in addition to their known branch-promoting activity. Unexpectedly, these VCA domains also destabilize existing branches, as well as SPIN90-Arp2/3 at filament pointed ends. Furthermore, cortactin and GMF, which respectively stabilize and destabilize Arp2/3 at branch junctions, have a similar impact on SPIN90-activated Arp2/3. However, unlike branch junctions, SPIN90-Arp2/3 at the pointed end of linear filaments is not destabilized by piconewton forces, and does not become less stable with time. It thus appears that linear and branched Arp2/3-generated filaments respond similarly to regulatory proteins, albeit with quantitative differences, and that they differ greatly in their responses to aging and to mechanical stress. These results indicate that SPIN90- and VCA-activated Arp2/3 complexes adopt similar yet non-identical conformations, and that their turnover in cells may be regulated differently.

## Introduction

Branched actin filament networks, such as the ones found in the lamellipodia of migrating cells, or in endocytic patches, are a hallmark example of dynamic actin networks in cells. Branches result from the activation of the Arp2/3 complex by the VCA domains of nucleation-promoting factors (NPFs), such as WASP or WAVE (See (Gautreau et al., 2022) for a review). The VCA domain binds to the Arp2/3 complex via its CA domain, and recruits an actin monomer (G-actin) via its V-domain (also known as WH2 domain). Two VCA domains can bind to the same Arp2/3 complex, and activate it more efficiently (Padrick et al., 2011; Zimmet et al., 2020). Following the binding of this large complex to the side of a pre-existing filament, commonly called the ‘mother’ filament, the VCA domains detach, thereby allowing the elongation of the branch (Smith et al., 2013). Actin subunits are added at the dynamic ‘barbed’ end of the branchs’, while its less dynamic ‘pointed’ end is stabilized and connected to the mother filament by the Arp2/3 complex.

The Arp2/3 complex can also be activated by the protein WISH/DIP/SPIN90, thereafter referred to as SPIN90 (Wagner et al., 2013). The binding of SPIN90 to the Arp2/3 complex results in the nucleation of an actin filament that elongates freely at the barbed end, while SPIN90-Arp2/3 remains attached to the pointed end. This activation mechanism generates de novo linear filaments, in the absence of a pre-existing mother filament.

The binding site of SPIN90 on Arp2/3 does not overlap with the binding sites of VCA but sits in the region where the mother filament would bind, in the VCA-induced branching reaction (Luan et al., 2018). Consistent with this, dip1 and wsp1, the fission yeast homologs of SPIN90 and WASP, respectively, can simultaneously bind and co-activate the Arp2/3 complex, to generate linear filaments (Balzer et al., 2020).

During endocytosis in yeast, linear SPIN90-nucleated filaments provide the initial mother filaments that will ‘prime’ the formation of branched networks (Balzer et al., 2020, 2019; Basu and Chang, 2011). In contrast, in the cortex of animal cells, SPIN90-induced nucleation decreases the branching density and favors the rapid elongation of filaments by formin mDia1 (Cao et al., 2020). It thus appears that, depending on the context, the two activation machineries of the Arp2/3 complex can synergize or compete. Whether they can be regulated independently is an open question: can one activation machinery be stimulated or inhibited without similarly affecting the other?

It was also shown that formin mDia1 could bind to the SPIN90-activated Arp2/3, indicating that it differs from the Arp2/3 complex in nascent branches (Cao et al., 2020). The high-resolution cryo-EM structures of Arp2/3 bound to SPIN90 (dip1 from yeast), and of Arp2/3 at branch junctions in cells, indicate that Arp2/3 undergoes similar conformation changes in both activation mechanisms (Fäßler et al., 2020; Shaaban et al., 2020). Small differences in inter-subunit contacts could also be detected (Fäßler et al., 2020). Understanding whether these conformational differences give rise to different responses to regulatory factors is a fundamental question that requires further investigation.

An important aspect of Arp2/3 regulation is the control of the stability of Arp2/3-generated filaments. With time, branches detach from mother filaments. Debranching is favored by the hydrolysis of ATP within the Arp2/3 complex (Dayel and Mullins, 2004; Le Clainche et al., 2003). Debranching can be modulated by proteins interacting with Arp2/3: branch junctions are stabilized by cortactin, and destabilized by GMF (Gandhi et al., 2010; Weaver et al., 2001). In addition, debranching is accelerated by mechanical tension applied to the branch (Pandit et al., 2020). The impact of these factors on SPIN90-activated Arp2/3 is an open question. More generally, the stability of SPIN90-Arp2/3 at the pointed end, which is important to understand the reorganization and disassembly of actin filament networks, remains to be established.

Here, we address these questions using purified proteins in single-filament experiments, using microfluidics and TIRF microscopy. We show that Arp2/3 nucleation is similarly stabilized by cortactin and destabilized by GMF, when activated either by VCA or SPIN90 to generate branches or linear filaments, respectively. Yet key differences, in particular regarding their sensitivity to mechanical stress and to Arp2/3 aging, indicate that the Arp2/3 complex adopts different conformations when activated by these two mechanisms.

## Results

Experiments were performed at 25°C, using actin from rabbit skeletal muscle, Arp2/3 from bovine brain, and recombinant mammalian isoforms of other proteins (see Methods)

### VCA enhances the nucleation of linear filaments by SPIN90-Arp2/3

We first sought to verify that, as with their yeast homologs (Balzer et al., 2020), mammalian VCA and SPIN90 can co-activate the Arp2/3 complex to generate linear filaments. To do so, we exposed surface-anchored SPIN90 to the Arp2/3 complex, and subsequently flowed in solutions of profilin:actin complex and different concentrations of VCA into the microfluidics chamber. Since the outcome of these experiments is very sensitive to small variations in protein concentration, we have taken advantage of a microfluidics assay to simultaneously perform experiments and control experiments, in two different regions of the same microchamber to minimize variations between experiments (see Methods, Fig 1A)(Jégou et al., 2011; Wioland et al., 2022, 2020).

**Figure 1.**
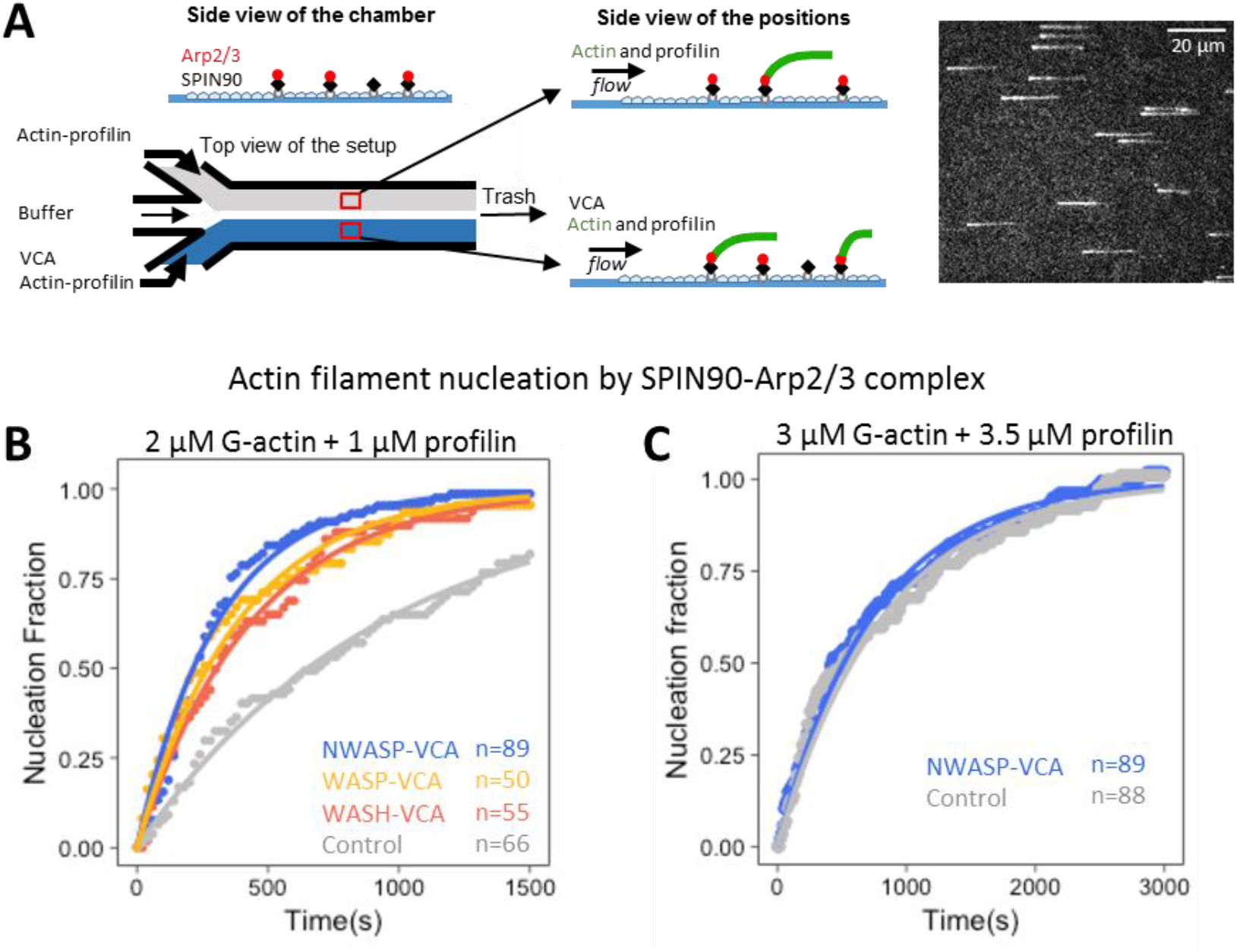
VCA domains accelerate the nucleation of filaments by SPIN90-Arp2/3. **A**. Schematic of the nucleation experiment. In a microfluidics chamber, SPIN90-Arp2/3 is attached to the coverslip surface, via the His-tag on SPIN90. Half of the chamber is then exposed to actin and profilin, while the other half is exposed to actin, profilin and VCA. Actin is labeled with Alexa488 (15%), and one can monitor filaments as they nucleate and grow from the surface. Right: TIRF microscopy image of actin filaments. **B**. Normalized number of filaments nucleated over time, from SPIN90-Arp2/3 exposed to 2 μM G-actin (15% labeled with Alexa488) and 1 μM profilin, with 0 or 0.5µM of GST-VCA from different NPFs. Solid lines are exponential fits, yielding nucleation rates k_nuc_=(1.06±0.03)×10^−3^ s^-1^ without VCA, and k_nuc_=(3.23±0.08)×10^−3^, (2.58±0.05)×10^−3^, and (2.28±0.03)×10^−3^ s^-1^ with VCA from N-WASP, WASP and WASH, respectively. **C**. Same as in (B), with 3 µM G-actin and 3.5 µM profilin.The exponential fits yield k_nuc_=(1.2±0.02)×10^−3^ s^-1^ without VCA, and k_nuc_=(1.3±0.02)×10^−3^ s^-1^ with 0.5µM GST-VCA from N-WASP. In (B,C), indicated values of n are the number of filaments observed in each individual experiment. These experiments were repeated three times, with similar results.

We found that the presence of VCA domains from either WASP, N-WASP or WASH, all induce a faster nucleation of filaments from SPIN90-Arp2/3 (Fig. 1B). VCA from N-WASP appeared to be slightly more potent than VCA from WASP and WASH in enhancing nucleation from SPIN90-Arp2/3 (with 2 µM G-actin and 1µM profilin, the estimated nucleation rate is 25-40% higher). We wondered if the VCA domains from these three NPFs ranked similarly for the activation of Arp2/3-induced branching. We thus compared the branch densities on pre-formed filaments, 90 seconds after exposing them for 90 seconds to Arp2/3, G-actin, and VCA from the different NPFs (see Methods). We observed the same ranking, with a more pronounced difference between NPFs: the branch density was 0.95, 0.43 and 0.27 branches/µm with VCA domains from N-WASP, WASP and WASH, respectively.

VCA recruits G-actin as part of the branched nucleation process (Chereau et al., 2005). With yeast homologs, it was found that the co-activation of Arp2/3 by VCA to form linear filaments required the recruitment of G-actin by VCA (Balzer et al., 2020). We thus hypothesized that the availability of G-actin would impact the ability of VCA to enhance the nucleation of filaments from SPIN90-Arp2/3. To test this hypothesis, we took advantage of profilin, which competes with VCA for G-actin. We found that, in the presence of an excess of profilin, VCA from N-WASP no longer enhanced the nucleation of filaments from SPIN90-Arp2/3 (Fig. 1C). This observation confirms that VCA enhances SPIN90-induced nucleation by recruiting actin monomers.

### VCA domains destabilize Arp2/3 at the pointed end of branches and linear filaments

During the course of our experiments, we also noticed that the nucleated filaments detached faster from the surface in the presence of VCA (Supp Fig S1). This indicates that VCA destabilizes SPIN90-Arp2/3 at pointed ends. To obtain further insights into this destabilization mechanism, we decided to quantify VCA-mediated detachment of SPIN90-Arp2/3-nucleated filaments, in the presence of 0.15 µM G-actin (to maintain filaments at a constant length).

Taking advantage of the sequential exposure provided by microfluidics, we elongated filaments from surface-anchored SPIN90-Arp2/3, and subsequently monitored their detachment, over time, as we exposed them to different concentrations of VCA domains (Fig. 2A-C). We found that all VCA domains accelerated the detachment of filaments from the surface, in a dose-dependent manner, albeit to different extents. Interestingly, the NPFs did not rank the same as for the enhancement of nucleation: VCA from N-WASP was still the most effective (increasing the detachment rate up to 16-fold in the 0-1 µM range) but WASP was much less effective than WASH.

**Figure 2.**
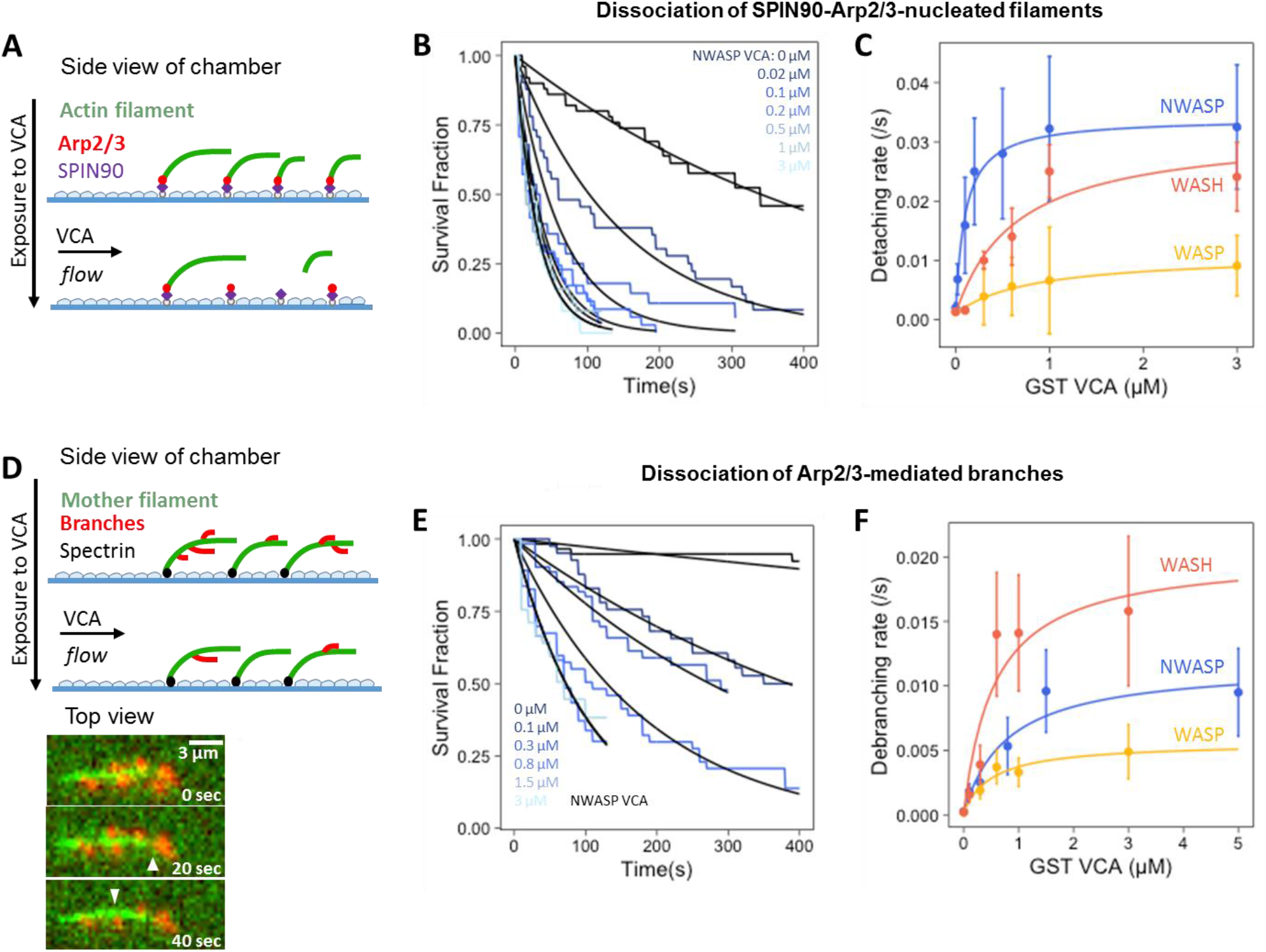
VCA domains accelerate the dissociation of both SPIN90-Arp2/3-nucleated actin filaments and Arp2/3-mediated branches. **(A-C)** Dissociation of SPIN90-Arp2/3-nucleated filaments. **(D-E)** Dissociation of Arp2/3-mediated branches. **A**. Sketch of the detachment experiment. Actin filaments were first nucleated from a SPIN90-Arp2/3-decorated surface, and elongated. Buffer with different concentrations of VCA was flowed in, and filament detachment events were monitored over time. **B**. The fraction of filaments still attached to the surface, versus time, for different concentrations of GST-NWASP-VCA. Black lines are exponential fits. **C**. Detachment rates, determined by exponential fits of survival curves (as in B) as a function of the concentration of VCA domains from different NPFs. The data are fitted by a Michaelis-Menten equation (resulting in an apparent K_D_ of 0.10±0.01, 0.65±0.36 and 0.82±0.14 µM, for N-WASP, WASH and WASP, respectively). **D**. Sketch of the debranching experiment. Alexa488(15%)-actin filaments (green) were elongated from surface-anchored spectrin-actin seeds, then exposed to VCA, Arp2/3 and Alexa568(15%)-actin to form branches (red). After 5 minutes, buffer with different concentrations of VCA was flowed in, and debranching events were monitored over time. Bottom: fluorescence microscopy images, showing debranching event (white arrowheads). **E**. The fraction of remaining branches, versus time, for different concentrations of GST-NWASP-VCA. Black lines are exponential fits. **F**. Debranching rates, determined by exponential fits of survival curves (as in E) as a function of the concentration of VCA domains from different NPFs. The data are fitted by a Michaelis-Menten equation (resulting in an apparent K_D_ of 0.57±0.35, 0.81±0.36 and 0.53±0.23 µM, for WASH, N-WASP and WASP, respectively). Error bars in (C,F) result from fits of the extremes of the 95% confidence intervals of survival functions (not shown in (B, E) for readability).

As dimerization enhances the branching activity of VCA domains (Padrick et al., 2008) we decided to compare the effectiveness of dimerized GST- and monomeric GFP-tagged VCA N-WASP. We found that dimerization had no impact on the ability of VCA to accelerate the nucleation of filaments by SPIN90-Arp2/3 (Supp Fig S2A). It did, however, enhance its ability to destabilize SPIN90-Arp2/3 at the pointed end of the filaments: the apparent K_D_ of GFP-VCA was 5.6-fold higher than that of GST-VCA (Supp Fig S2B-D).

To determine if VCA domains could also destabilize Arp2/3-mediated branches, we also performed similar experiments on branched filaments (Fig. 2D-F). We found that all VCA domains accelerated debranching, in a dose-dependent manner. However, the NPFs did not rank the same as for the detachment of SPIN90-Arp2/3-nucleated filaments: VCA from WASP was still the least effective, but VCA from WASH was the most effective (accelerating debranching up to 83-fold in the 0-3 µM range). Intriguingly, this ranking also differed from the branching situation, where we found VCA from WASH to be the least effective.

Since the recruitment of G-actin by VCA is necessary both for branching and for the enhancement of SPIN90-induced nucleation (Balzer et al., 2020; Chereau et al., 2005)(Fig. 1C), we wondered how the availability of G-actin would affect the ability of VCA to induce debranching and/or destabilize SPIN90-Arp2/3 at filament pointed ends. We thus repeated the experiments of figure 2 with different amounts of G-actin. We found that the ability of VCA to destabilize Arp2/3, either at branch junctions or at the pointed end of linear filaments, was reduced by the presence of G-actin (Supp Fig S3). These observations suggest that the ability of VCA to destabilize Arp2/3-nucleated filaments relies on the availability of its V-domain.

### The debranching factor GMF destabilizes SPIN90-Arp2/3-nucleated filaments

Given our observations, we wondered whether branch destabilizers such as GMF also impact the stability of SPIN90-Arp2/3-nucleated filaments. To do so, we performed detachment experiments similar to the ones in Figure 2A, where we exposed the filaments to different concentrations of GMF. We found that GMF accelerated the detachment of filaments from the surface (Fig 3), with a higher affinity than VCA (Fig 2A-C): K_D_=25+/-130 nM for GMF, compared to 100-820 nM for the VCA domains from different NPFs.

**Figure 3.**
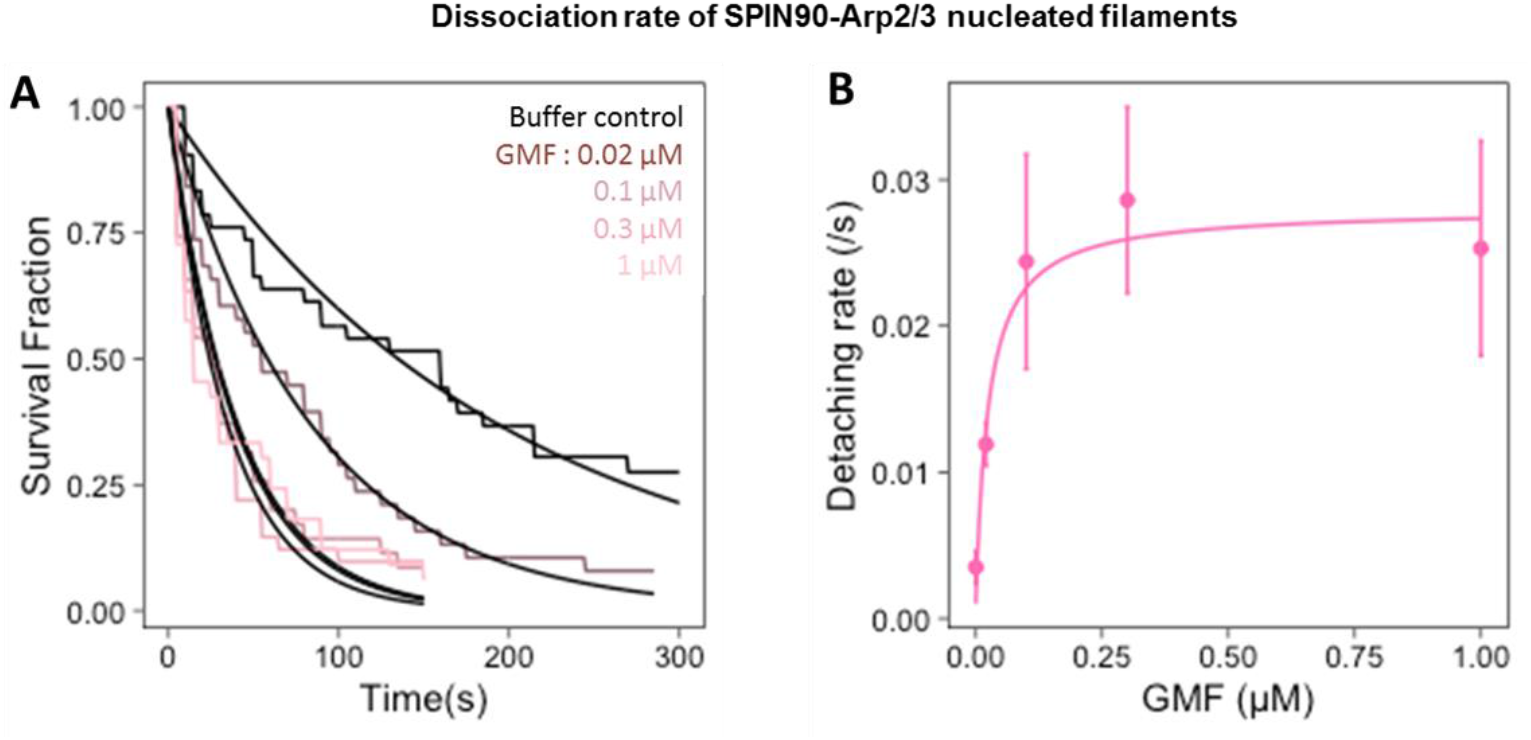
GMF accelerates the detachment of SPIN90-Arp2/3-nucleated actin filaments. **A**. The fraction of filaments still attached to the surface by SPIN90-Arp2/3, over time, when exposed to different concentrations of GMF. For each experiment, n=33-42 filaments. Black lines are exponential fits. **B**. Detachment rates, determined by exponential fits of survival curves shown in A, as a function of the concentration of GMF. The error bars are determined by the exponential fits in A. The data are fitted by a Michaelis-Menten equation, yielding an apparent dissociation constant K_D_=25±11 nM.

### VCA and GMF generate free pointed ends as they dissociate the Arp2/3 complex from both SPIN90 and the filament

So far, our readout of the destabilization of SPIN90-Arp2/3 at the filament pointed end is the detachment of the filament from the surface. Since SPIN90 is strongly anchored to the surface via their His-tag (see Methods), the detachment of the filament could result either from the rupture of the SPIN90-Arp2/3 bond, or from the removal of the Arp2/3 complex from the filament pointed end.

To determine whether the SPIN90-Arp2/3 interaction is destabilized by GMF and VCA, we examined whether the Arp2/3 complex was still bound to SPIN90 after the detachment of the actin filament, in our microfluidics assay (Fig. 4A-C). To do so, we quantified the fraction of functional SPIN90 locations that were able to renucleate a filament, following the departure of the initial actin filament, in different conditions. The new filaments took several tens of seconds to appear, consistent with renucleation, and ruling out regrowth from broken filaments which would be instantaneous. Based on the density of nucleated filaments in our experiments, it is very unlikely that renucleated filaments result from a different, co-localized SPIN90.

**Figure 4.**
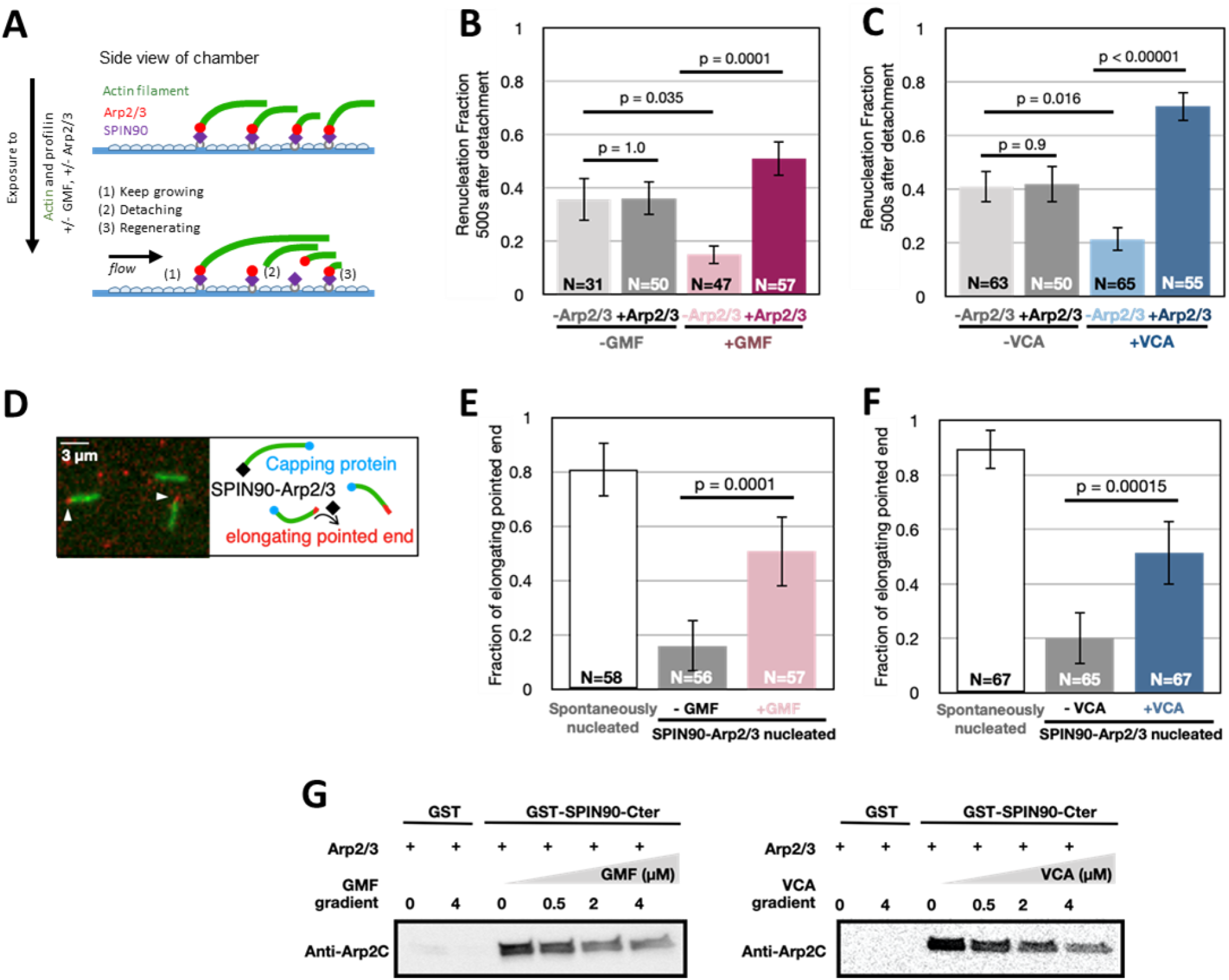
GMF and VCA accelerate the detachment of Arp2/3 from both SPIN90 and filament pointed ends. **A**. Sketch of the renucleation assay, using microfluidics. The SPIN90-Arp2/3 complex-decorated surface is exposed to 2 μM (15% Alexa488)G-actin and 1 μM profilin, with 0 or 0.5 μM GMF, and with 0 or 5 nM Arp2/3. Filaments nucleated by SPIN90-Arp2/3 elongate (1) and, over time, detach from the surface (2). Some of them renucleate, from the same location (3). **B-C**. The fraction of SPIN90 that renucleated a filament 500 seconds after the detachment of the initial filament, was quantified for different conditions The error bars represent the 95% confidence intervals. P-values were calculated based on a chi-square test. **D**. Sketch and microscope image of the pointed end uncapping assay. Preformed filaments made with (15% Alexa488)-actin were mixed with 10 nM Capping protein, 1 μM (15% Alexa568)-G-actin, and 0 or 0.5 μM GMF (or 0.5 µM VCA). In the microscope image (left), filaments were nucleated by SPIN90-Arp2/3, and exposed to 0.5 µM GMF. Slow elongation of the filaments at one end (white arrowheads) indicates freely growing pointed ends. **E-F**. The fraction of elongating, i.e. not capped, pointed ends was determined for different conditions. The error bars represent the 95% confidence intervals. P-values were calculated based on a chi-square test. The experiments in (A-F) were repeated twice, with similar results. **G**. GST pull-down assays showing that GMF and VCA interfere with Arp2/3 binding SPIN90. GST beads, decorated with GST or GST-SPIN90-Cter (the functional domain of SPIN90), were incubated with 0.4 μM Arp2/3 and gradient concentrations of GMF or VCA for 1 hour at room temperature. After the unbound protein was washed out, the amount of Arp2/3 attached to the beads was detected by Anti-2C antibody. We verified that the beads were loaded with equal amounts of GST or GST-SPIN90 (Supp Fig S4).

When only actin and profilin were flowed into the chamber, we observed that 36% of surface-anchored SPIN90 were able to renucleate a filament, within the 500 seconds following the detachment of their first filament (Fig. 4A-C). Based on our earlier experiments (Fig. 1B), this number is the expected nucleation fraction after that time. This confirms that the population of anchored-SPIN90 from which filaments were first nucleated is still attached to the surface. This also indicates that, upon detachment of the filament, the Arp2/3 complex stays bound to SPIN90. Consistent with this interpretation, supplying fresh Arp2/3 during the experiment did not increase the fraction of renucleated filaments (Fig. 4B-C).

In the presence of GMF, however, the renucleation fraction dropped to 15%, and was rescued when fresh Arp2/3 was supplied (Fig. 4B). This indicates that GMF destabilizes the SPIN90-Arp2/3 bond in the presence of actin. We obtained similar results with the VCA domain of N-WASP (from here onward, VCA refers to GST-NWASP-VCA) indicating that VCA also destabilizes the SPIN90-Arp2/3 bond in the presence of actin (Fig. 4C). However, these experiments do not tell us if the separation of Arp2/3 from SPIN90 causes filament detachment, or if it occurred after. They also do not indicate whether Arp2/3 remains attached to the pointed end of the departing filament.

To determine whether the Arp2/3 complex remained bound to the pointed ends of the filaments exposed to GMF or VCA, we performed experiments in open chambers (no microfluidic flow) where we could monitor the filaments’ ability to elongate from their pointed ends. To evaluate the fraction of capped pointed ends, we exposed preformed Alexa488(15%)-actin filaments to barbed end capping proteins, and to Alexa568(15%)-G-actin (Fig. 4D). The vast majority of spontaneously nucleated filaments were able to elongate from their pointed ends, validating the rationale of the assay (Fig. 4E-F). In contrast, most filaments nucleated by SPIN90-Arp2/3 were unable to do so, indicating that their pointed ends were capped by SPIN90-Arp2/3. In the presence of GMF or VCA, the fraction of elongating pointed ends increased significantly (Fig. 4E-F). This demonstrates that the Arp2/3 complex was removed from the pointed end following, or during, the destabilization of SPIN90-Arp2/3 by GMF or VCA.

To determine if SPIN90-Arp2/3 is also destabilized by GMF or VCA in the absence of a filament, we performed bead pull-down assays. We show that GMF and VCA both decrease the amount of Arp2/3 bound to SPIN90-decorated beads (Fig 4G). This observation indicates that, in the absence of actin, GMF and VCA interfere with the binding of Arp2/3 to SPIN90.

### Piconewton forces promote debranching, but have no impact on SPIN90-nucleated filaments

It was recently shown that debranching is accelerated when (sub)piconewton forces are applied to the branch (Pandit et al., 2020). We therefore wondered whether tensile forces also destabilize SPIN90-Arp2/3 at the pointed end of filaments.

By controlling the flow rate in our microfluidics assay, we can vary the force applied to actin filaments (Jégou et al., 2013). Forces ranging from 0.2 to 2 pN accelerated debranching (Fig. 5A). This observation remained true in the presence of VCA and GMF: pulling forces further accelerated debranching (Fig 5 B-D). In contrast, we found that forces up to 2 pN had no impact on the detachment of SPIN90-Arp2/3-nucleated filaments, even in the presence of VCA or GMF (Fig. 5E-H).

**Figure 5.**
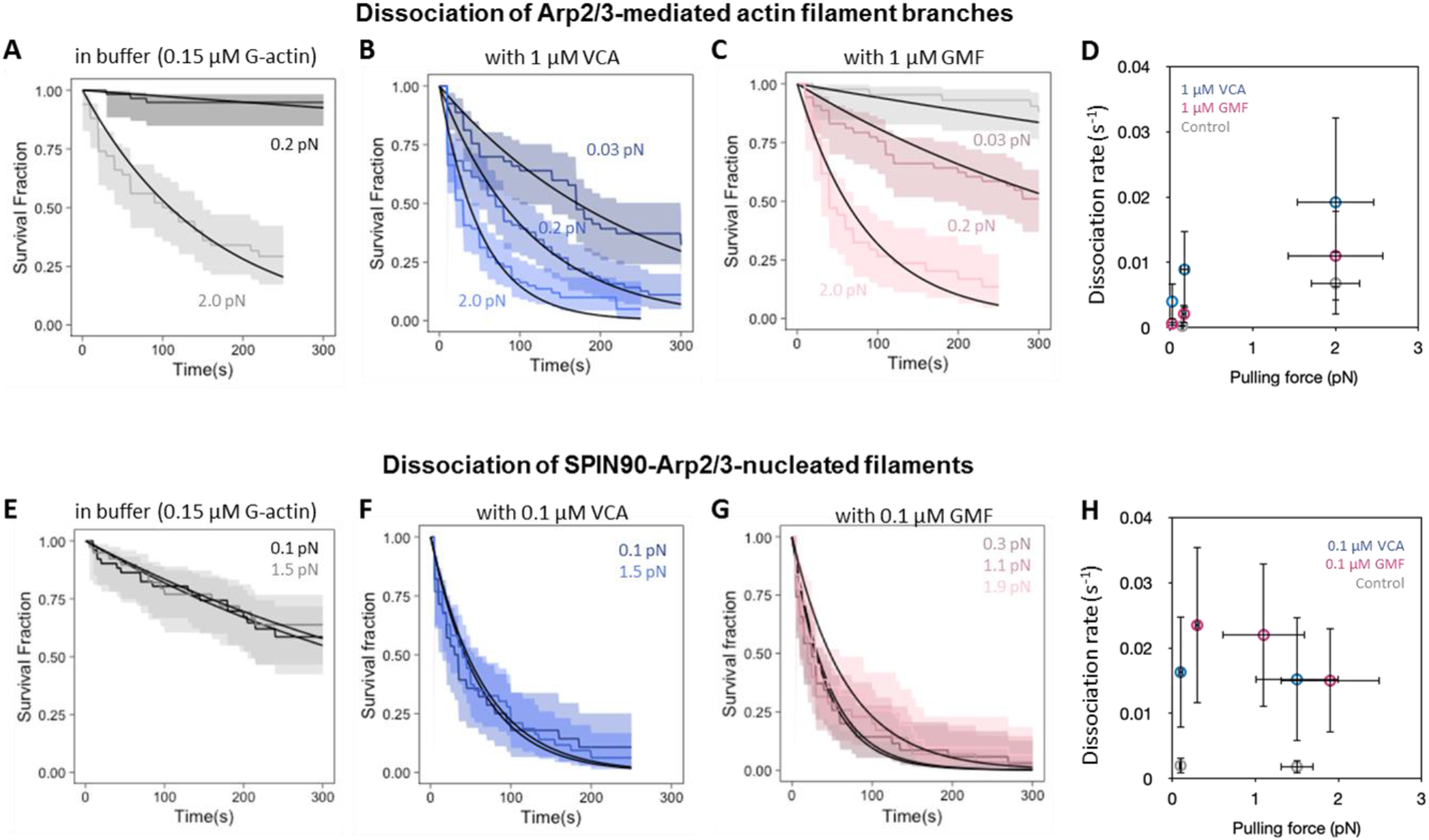
Impact of mechanical tension on debranching and on the detachment of SPIN90-Arp2/3-nucleated filaments. **A-C**. Survival fractions for branches exposed to different forces in the microfluidics chamber, while being exposed to 0.15µM G-actin, alone (gray curves, A), with 1 µM VCA (blue curves, B), or with 1 µM GMF (red curves, C). The shaded areas represent 95% confidence intervals. Black lines are exponential fits. **D**. Summary of the debranching rates, resulting from the exponential fits in (A-C). **E-G**. Survival fractions for SPIN90-Arp2/3-nucleated filaments exposed to different forces in the microfluidics chamber, while being exposed to 0.15µM G-actin, alone (gray curves, E), with 0.1 µM VCA (blue curves, F), or with 0.1 µM GMF (red curves, G). The shaded areas represent 95% confidence intervals. Black lines are exponential fits. **H**. Summary of the detachment rates for SPIN90-Arp2/3-nucleated filaments, resulting from the exponential fits in (E-G).

To make sure that this result does not depend on our anchoring SPIN90 by its C-terminus (see Methods), we also attached the protein by its N-terminus (Supp Fig S5A). Again, we found that mechanical tension had no impact on filament detachment. We also verified that tension had no impact on detachment when exposing the filaments to GFP-VCA instead of GST-VCA (Supp Fig S5A), or to a higher concentration (1 µM) of GST-VCA (Supp Fig S5B).

### Aging promotes debranching but has no impact on SPIN90-nucleated filaments

Debranching is enhanced as ATP is hydrolyzed in Arp2 and Arp3, within the Arp2/3 complex at a branch junction, within minutes (Le Clainche et al., 2003; Pandit et al., 2020). Branches thus become less stable over time. Quantifying ATP hydrolysis in SPIN90-activated Arp2/3 is particularly challenging, because of the large background of ATP hydrolysis taking place in F-actin, and because SPIN90 activates even lower amounts of Arp2/3 than the branching reaction. Instead, to gain insights into this question, we sought to determine if time destabilizes SPIN90-Arp2/3 at filament pointed ends, as seen with branch junctions.

In our experiments up to now, we observed the detachment of branches and SPIN90-nucleated filaments on average 5 minutes following their nucleation. Here, we aged filaments and branches for an additional delay while exposing them to minimal forces with a gentle flow, and subsequently monitored their detachment, on average 25 minutes after their nucleation.

We confirmed that 25-minute-old branches detach faster than 5-minute-old branches, and found that this remains true when debranching is enhanced by VCA (Fig. 6A). In contrast, we observed no significant difference between 5- and 25-minute-old SPIN90-Arp2/3-nucleated filaments, with or without VCA (Fig. 6B).

**Figure 6.**
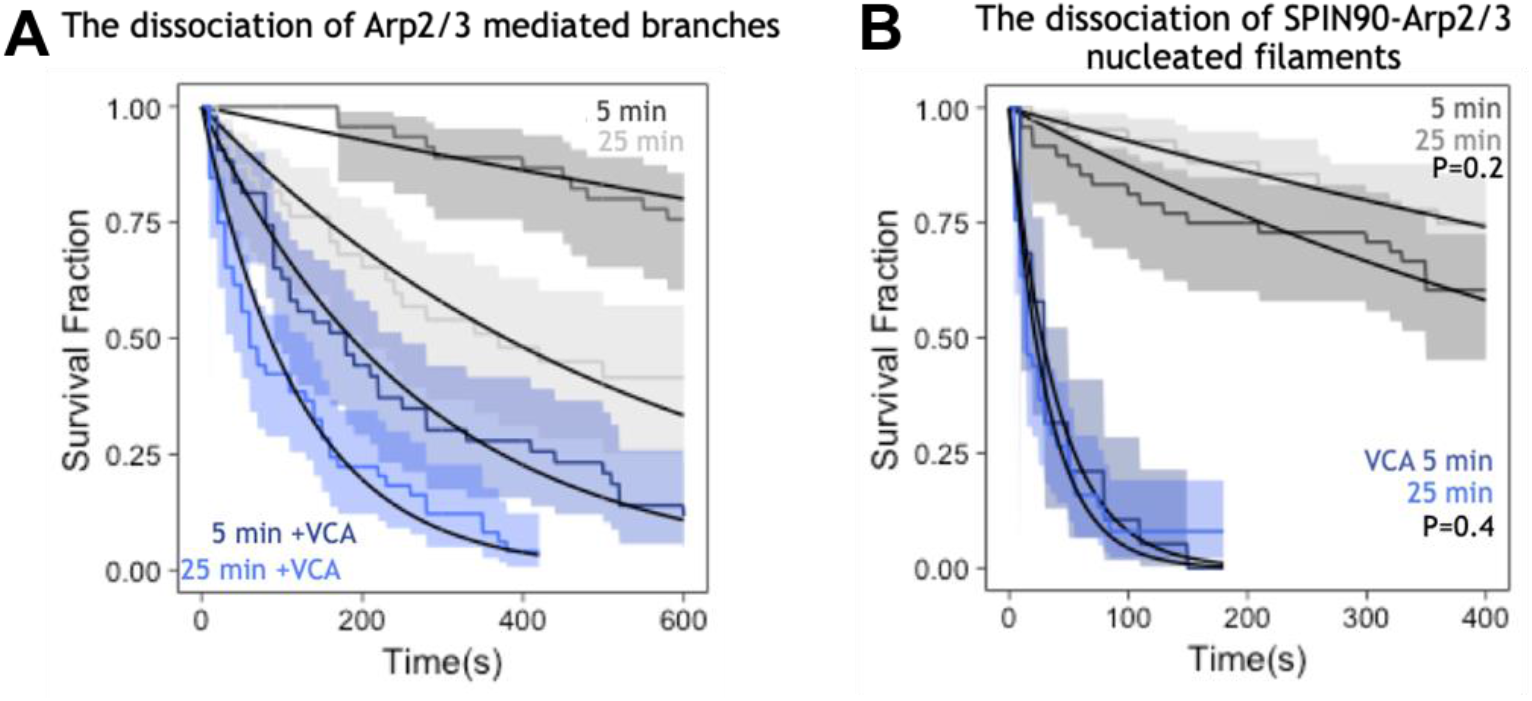
Aging destabilizes branches but has no measurable effect on SPIN90-Arp2/3-nucleated filaments. **A**. Fraction of remaining branches, over time, starting at an average of 5 and 25 minutes after branch nucleation, in the presence (blue) or absence (gray) of VCA. Lines are exponential fits, yielding debranching rates of (3.7±0.6)×10^−4^ s^-1^ and (20±8)×10^−4^ s^-1^ without VCA for 5- and 25-minute-old branches, respectively; and debranching rates of (3.8±1.3)×10^−3^ s^-1^ and (8.2±2.8)×10^− 3^ s^-1^ with 0.5 µM VCA for 5- and 25-minute-old branches, respectively. **B**. Fraction of attached SPIN90-nucleated filaments, over time, starting at an average of 5 and 25 minutes after branch nucleation, in the presence (blue) or absence (gray) of VCA. Lines are exponential fits, yielding detachment rates of (12±7)×10^−4^ s^-1^ and (7±3)×10^−4^ s^-1^ without VCA for 5- and 25-minute-old filaments, respectively (p=0.2, logrank test); and detachment rates of (2.5±1.1)×10^−2^ s^-1^ and (3.1±1.7)×10^−2^ s^-1^ with 1 µM VCA for 5- and 25-minute-old filaments, respectively (p=0.4, logrank test). Shaded areas represent 95% confidence intervals. The flowing solutions apply a weak force to the filaments (< 1pN) and branches (0.2 pN).

### Cortactin stabilizes branches and SPIN90-Arp2/3 at filament pointed ends

Finally, we asked if cortactin, a protein proposed to interact with the Arp2/3 complex to stabilize branches (Abella et al., 2016; Weaver et al., 2001), could have a similar effect on SPIN90-Arp2/3-nucleated filaments. Since the stabilizing activity of cortactin was never observed directly on filament branches, we sought to confirm this property by carrying out debranching experiments as in Figure 2D. We observed that 30-100 nM cortactin indeed slows down debranching, by approximately 2-fold (Fig. 7A, Supp Fig. S6).

**Figure 7.**
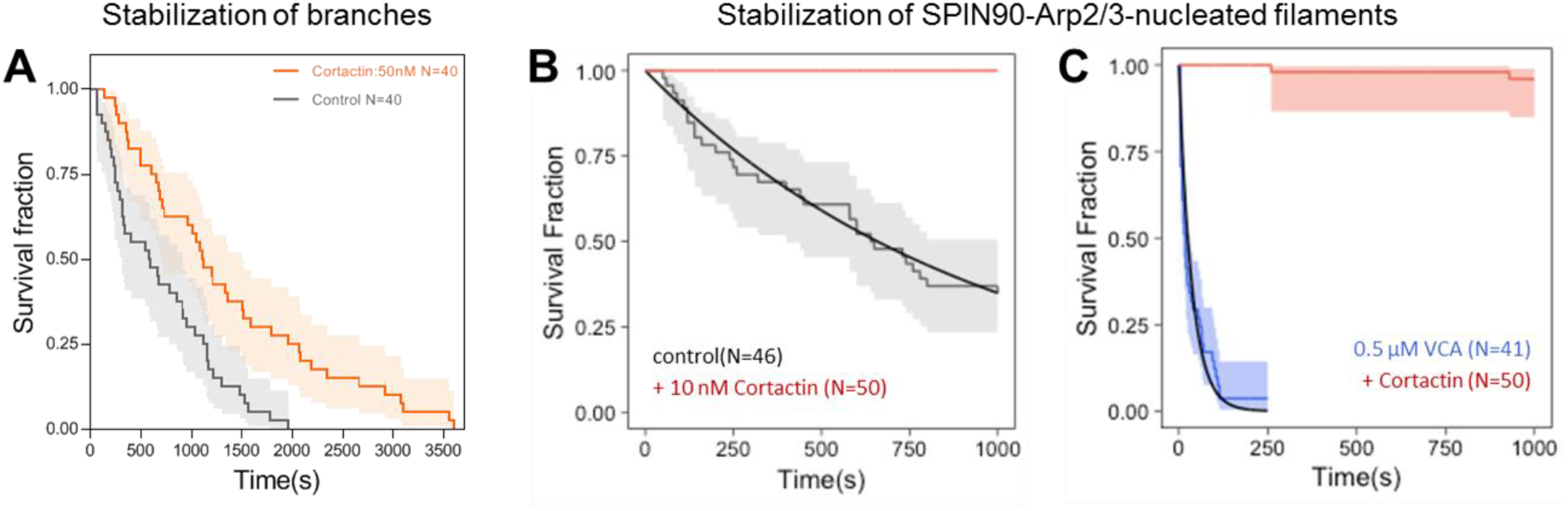
Cortactin stabilizes Arp2/3-mediated branches as well as SPIN90-Arp2/3 at the pointed end of filaments. **A**. The fraction of remaining branches, versus time, exposed to 0.16 µM G-actin (black) supplemented with 50 nM cortactin (red). The branch junctions were exposed to an average force of 1.4 pN. Similar results were obtained with different forces and different protein concentrations (Supp Fig S6). **B**. The fraction of remaining SPIN90-Arp2/3-nucleated filaments, versus time, exposed to 0.15 µM G-actin (black) supplemented with 10 nM cortactin (red). **C**. The fraction of remaining SPIN90-Arp2/3-nucleated filaments, versus time, exposed to 0.15 µM G-actin, with 0.5 µM VCA (black) supplemented with 10 nM cortactin (red). Shaded areas represent 95% confidence intervals.

We next monitored the nucleation and the detachment of actin filaments from a SPIN90-Arp2/3-decorated surface, in the presence of cortactin. We found that cortactin very efficiently stabilizes SPIN90-Arp2/3 at filament pointed ends (Fig. 7B): 10 nM cortactin were enough to prevent all filament dissociation from the surface over the course of the experiment (1000 seconds). We then wondered what would be the outcome of SPIN90-nucleated filaments in the presence of both cortactin and a destabilizer like VCA. We found that, in the presence of 0.5 µM VCA, the addition of 10 nM cortactin reduces the filament detachment rate by 3 orders of magnitude (Fig. 7C).

We also observed that cortactin accelerates the nucleation of filaments by SPIN90-Arp2/3, even in the presence of an excess of profilin (Supp Fig S7). The latter result is in contrast with VCA, whose enhancement of SPIN90-induced nucleation requires G-actin (Fig. 1). This observation is consistent with the fact that cortactin does not bind G-actin.

## Discussion

In cells, the Arp2/3 complex can be activated by VCA to generate branches, or by SPIN90 to generate linear filaments. Whether these two mechanisms can be regulated separately is an outstanding question. Here, we measured the impact of different regulatory factors on both mechanisms.

We found that both systems were stabilized by cortactin, and destabilized by GMF as well as by the VCA domains of NPFs. While these regulatory proteins similarly affect branched and linear Arp2/3-generated filaments, they do so with clear quantitative differences. For example, cortactin appears far more efficient at stabilizing SPIN90-Arp2/3 at the pointed end than at stabilizing branches (Fig. 7). Likewise, SPIN90-Arp2/3 at pointed ends is destabilized by lower concentrations of GMF, compared to branches (Fig. 5). Also, we found that VCA from different NPFs did not rank the same in their ability to destabilize SPIN90-Arp2/3 and branches (Fig. 2CF). These quantitative biochemical differences suggest that proteins in cells may differently regulate branched and linear Arp2/3-generated filaments, thereby providing a means to control the two Arp2/3 machineries independently.

Additional factors may further tune these regulatory activities. For instance, we found that aging and mechanical forces, which both strongly favor debranching, have little effect on the stability of SPIN90-Arp2/3 at pointed ends (Fig. 5 and 6). This is an important feature in the cellular context, where actin networks containing both types of Arp2/3-generated filaments, such as the cell cortex or during endocytosis, are exposed to mechanical stress. It indicates that mechanical stress can contribute to debranching, but that regulatory proteins are likely required to remove SPIN90 from the pointed ends.

The differences we observe, where it appears that SPIN90-Arp2/3 is globally more sensitive to regulatory proteins and less sensitive to aging and mechanical tension than branch junctions, can be linked to conformational differences between activated Arp2/3 in the two situations. The cryo-EM structures of the two forms of activated Arp2/3 exhibit small differences in contacts between subunits ArpC3 and Arp2, and between ArpC5/C5L and Arp2 (Fäßler et al., 2020; Shaaban et al., 2020). These differences are likely to affect interactions with GMF and VCA, which can both bind to Arp2 (Luan and Nolen, 2013; Zimmet et al., 2020). Branch junctions become less stable as ATP in Arp2 and Arp3 is hydrolyzed, within minutes after a branch is formed (Dayel and Mullins, 2004; Le Clainche et al., 2003). In contrast, ATP is still detected in Arp2 and Arp3 by cryo-EM, hours after activation by SPIN90 (Shaaban et al., 2020). This suggests that SPIN90-activated Arp2/3 does not hydrolyse ATP, or does so very slowly, and is consistent with our observation that SPIN90-Arp2/3 does not seem to age (Fig. 6). The structural basis for the mechanical robustness of SPIN90-Arp2/3 is less clear. SPIN90 binds primarily to ArpC4 while, in a branch junction, the mother filament forms an extensive interface with several subunits of Arp2/3 (Fäßler et al., 2020; Luan et al., 2018). One could thus have expected Arp2/3 to detach more readily from SPIN90 than from the mother filament. Our results suggest that yet undetected conformational differences may be at play.

Our observation that VCA enhances debranching (Fig. 2) indicates that VCA can bind to Arp2/3 at the branch junction. This was unexpected because VCA detaches from Arp2/3 on the side of the mother filament during the branching reaction (Smith et al., 2013). This result further suggests that VCA, beyond its role in the formation of branched filament networks, may also contribute to their reorganization by enhancing debranching. In cells, NPFs expose their VCA domains upon activation and binding to the membrane, meaning that branch junctions would need to contact the NPF-decorated membrane in order for debranching to be enhanced. This situation could be encountered during endocytosis, for example, where the branched filament network is densely packed against the membrane, and where debranching appears to be required to complete endocytic internalization (Martin et al., 2006). A limitation, however, would come from the high concentration of cytoplasmic profilin-actin, which would bind to the polyproline domain of NPFs and ultimately load G-actin onto their V-domain (Bieling et al., 2018). Unless G-actin is locally depeleted, this would likely inhibit, at least partially, the destabilizing activity of NPFs (Supp Fig S3).

In cells, where the turnover of actin networks is tightly controlled, the state of filament pointed ends is of primary importance. Early EM observations of branched filaments detected the Arp2/3 complex also at the pointed end of linear filaments, suggesting that Arp2/3 could act as a pointed end capper (Mullins et al., 1998). The persistence of SPIN90 with Arp2/3 at pointed ends makes this cap very stable, blocking filament depolymerization and renealling, and preventing SPIN90 from nucleating other filaments (Balzer et al., 2019). We show that the destabilization of SPIN90-Arp2/3 by GMF or VCA results in free pointed ends and renews the pool of free SPIN90 and Arp2/3 complexes (Fig. 4). In cells, we expect such mechanisms to contribute to controlling the state of pointed ends, and thus to controlling the stability and turnover of actin filament networks. Future studies should reveal if other inhibitors of branching, like Arpin or coronin, can play a similar role with SPIN90-Arp2/3 at pointed ends.

## METHODS

### Biochemistry

#### Protein preparation

Skeletal muscle actin was purified from rabbit muscle acetone powder following the protocol from (Spudich and Watt, 1971), as described in (Wioland et al., 2017).

We have recombinantly expressed the following proteins in bacteria, with a N-ter GST tag: human N-WASP-VCA (392-505, uniprot O00401), WASP-VCA (418-502, uniprot P42768), mouse WASH-VCA (319-475), human SPIN90 full-length (1-722, uniprot Q9NZQ3), SPIN90 Cter (267-722), and mouse cortactin full length (1-546 Q60598). Expression was performed in E.Coli Rosetta 2 (DE3) for 16h at 18 °C. GST-tagged proteins were purified by affinity chromatography over a Sepharose 4B GSH affinity column (Cytiva) in GST-buffer (20 mM Tris pH 7.5, 500 mM NaCl, 1 mM DTT). The GST protein was eluted with the same buffer supplemented with 50 mM GST. The purification is followed by gel filtration over a Hiload Superdex 200 column (Cytiva) in GF-buffer (20 mM Tris pH 7.5, 50 mM KCl, 1 mM DTT, 5% Glycerol). Before the gel filtration step, the GST-tag of GFP-N-WASP-VCA and cortactin was cut by PreScission Protease (Sigma-Aldrich) overnight at 4°C, to obtain monomeric N-WASP-VCA and untagged full length cortactin.

C-terminal His-tagged versions of recombinant human SPIN90 full-length (1-722) and drosophila GMF (1-138, uniprot Q9VJL6) were purified by affinity chromatography over a HisTrap HP column (Cytiva) in His-buffer (20 mM Tris pH 7.5, 500 mM NaCl, 1 mM DTT, 20 mM Imidazole) and eluted in the same buffer supplemented with 300 mM Imidazole. The purification was polished by gel filtration over a Hiload Superdex 200 column (Cytiva) in 20 mM Tris pH 7.5, 500 mM NaCl, 1 mM DTT.

Recombinant human profilin1 was expressed and purified, as described in (Cao et al., 2018). Arp2/3 was purified from bovine brain, as previously described (Le Clainche and Carlier, 2004).

#### Protein labelling

Actin was fluorescently labeled on the surface lysine 328, using Alexa-488 or Alexa-594 succinimidyl ester (Life Technologies) as described in detail in (Romet-Lemonne et al., 2018). To minimize potential artifacts caused by the presence of the fluorophores, we typically used an actin labeling fraction of 15 %.

### Microscopy

#### Buffers

All microfluidics experiments were performed in F-buffer: 5 mM Tris HCl pH 7.0, 50 mM KCl, 1 mM MgCl_2_, 0.2 mM EGTA, 0.2 mM ATP, 10 mM DTT, 1 mM DABCO and 0.1% BSA. Open chambers experiments (figure 4D-F) were performed using the same buffer supplemented with 0.3% methylcellulose 4000 cP.

#### Single filament assays in microfluidics

Microfluidics experiments were done with Poly-Dimethyl-Siloxane (PDMS, Sylgard) chambers based on the original protocol from (Jégou et al., 2011). The chambers have cross-shaped channels with three inlets and one outlet. A MFCS system and Flow Units (Fluigent) were used to control and monitor the microfluidic flows. PDMS chambers were cleaned and mounted by following the protocol described in (Cao et al., 2018). The experiments were performed on a Nikon TiE inverted microscope, equipped with a 60x oil-immersion objective, in F-buffer. The temperature was controlled and set to 25°C (using an objective heater from Oko-lab), except for the experiments with GMF where the temperature was a few degrees above 25°C. Actin filaments were visualized using TIRF microscopy (iLAS2, Gataca Systems) with 150 mW tunable lasers. Images were acquired using an Evolve EMCCD camera (Photometrics), controlled with the Metamorph software (version 7.10.4, from Molecular Devices).

#### SPIN90-Arp2/3 assays

Full length GST-SPIN90 or SPIN90-His were specifically anchored to the coverslip surface of a microfluidics chamber using specific antibodies adsorbed to the surface, prior to passivation by exposing the chamber to a 3% BSA solution for at least 5 minutes, as described in (Cao et al., 2020). The chamber was subsequently exposed to 30nM Arp2/3 complex for 5 minutes and rinsed with F-buffer.

- **SPIN90-Arp2/3 filament nucleation** The nucleation rate is measured by exposing surface-anchored His-tagged SPIN90-Arp2/3 to 2 μM 15% Alexa-488 actin and 1 μM profilin, with or without 0.5 μM of the different VCA domains (figure 1), or in the presence or absence of 50 nM cortactin (Supp Fig S7).
- **Dissociation of SPIN90-Arp2/3-nucleated filaments** Surface-anchored SPIN90-Arp2/3 was exposed to 0.7 μM 15% Alexa-488 actin for 5 minutes to nucleate and elongate actin filaments, in the presence of varying concentrations of the different VCA domains (figure 2A-C). The survival fractions of SPIN90-Arp2/3-nucleated filaments over time were fitted with a single exponential function that yields the dissociation rates of filament from SPIN90-Arp2/3 for each condition. The dissociation rates were plotted versus the concentration of VCA, and a fit using the Michaelis-Menten equation was used to obtain the affinity constant and maximum dissociation rate. The dissociation rate of SPIN90-Arp2/3 nucleated actin in the presence of cortactin was measured following the same procedure (Fig 7). The dissociation rate of SPIN90-Arp2/3 nucleated actin in the presence of GMF was measured following the same procedure but using the full-length GST-SPIN90 construct (Fig 3). To measure the impact of time on the dissociation rate, generated actin filaments were incubated with 0.15 μM G-actin for an extra duration of 20 minutes, before being exposed to buffer only or 0.5 µM of N-WASP-VCA (Fig 6).
- **Impact of mechanical tension on the detachment of SPIN90-Arp2/3-nucleated filaments**. With the same experimental procedure described in the previous paragraph, various pulling forces can be applied to the filaments by varying the flow rate and filament length (Jégou et al., 2013) (Fig 5).
- **Detachment of Arp2/3 from both SPIN90 and the filament pointed end (Figure 4)**. To assess the ability of GMF to separate Arp2/3 from SPIN90, surface-anchored GST-SPIN90-Arp2/3-nucleated filaments were exposed to 2 μM 15% Alexa-488 labeled G-actin, 1 μM profilin, with and without 0.5 μM GMF, and with or without 5 nM Arp2/3 (figure 4B). The renucleation fraction of a randomly picked population of SPIN90-Arp2/3-nucleated filaments was assessed after 500 seconds. To assess the ability of GST-N-WASP-VCA to separate Arp2/3 from SPIN90, surface-anchored SPIN90-His-Arp2/3-nucleated filaments were alternatively exposed to 0.5 μM GST-N-WASP-VCA or buffer for 30 seconds, then to a nucleation solution containing 2 μM 15% Alexa-488 labeled actin and 1 μM profilin, with or without 5 nM Arp2/3 for 90 seconds. In this assay, VCA and Arp2/3 were not mixed together with actin to avoid unnecessary generation of branched actin in solution and on the surface-anchored filaments. The renucleation fraction of a randomly picked population of SPIN90-Arp2/3-nucleated filaments was assessed after 500 seconds. (Fig 4C).
- **Single filaments in open chambers (Figure 4D-F)** Experiments aimed at observing pointed end elongation were not performed using microfluidics, but using so-called ‘open chambers’, i.e. mPEG-silane-passivated flow chambers made following the protocol described in (Cao et al., 2020). 50 nM Arp2/3, 1 μM SPIN90 Cter, 0.5 μM 15% Alexa-488 actin were mixed together for 3 minutes to generate SPIN90-Arp2/3 nucleated actin filaments. As a positive control, 1 μM 15% Alexa-488 actin were mixed together for 3 minutes to generate spontaneously nucleated actin filaments. 4 μL of SPIN90-Arp2/3 nucleated filaments or 2 μL of spontaneously nucleated actin filaments were mixed with 10 nM Capping protein, 1 μM 15% Alexa-568 actin with or without 0.5 μM GMF and flowed into the chamber immediately, using an F-buffer supplemented with 0.3% methylcellulose 4000 cP. The fraction of actin filaments with an elongating pointed end is quantified at time t=350 seconds after solution mixing. To study the uncapping effect of VCA, the same experiment was done except that we used 5 nM Arp2/3, 2 μM SPIN90 Cter, 0.5 μM 15% Alexa-488 actin to generate SPIN90-Arp2/3 nucleated actin filaments to minimize the amount of Arp2/3 in the system.

#### Arp2/3-mediated branches assays

- **Dissociation of branches by VCA domains, cortactin, and forces** The coverslip surface of a microfluidics chamber was functionalized by exposing it to 2.5 pM spectrin-actin seeds for 5 minutes, followed by a passivation with a solution containing 3% BSA for 5 minutes. Mother filaments were grown from seeds by exposing them to 0.7 μM 15% Alexa-488 actin. Mother filaments were then exposed to 0.15 μM unlabelled actin for 1 minute. To generate the branches, the mother filaments are sequentially exposed to a solution of 30 nM Arp2/3 and 0.25 μM VCA for exactly 2 minutes, followed by a solution of 0.5 μM 15% Alexa-568 actin for 5 minutes in order to obtain branches in a different color than the one of the mother filaments. Arp2/3-generated branches were exposed to 0.15 μM 15% Alexa-488 actin in the presence or absence of varying concentrations of the different VCA domains (Fig 2D-F) or 50 nM cortactin (Fig 7A). The survival fractions of branches over time were fitted with a single exponential function that yields the dissociation rates of branches for each condition. The dissociation rates were plotted versus the concentration of VCA, and a fit using the Michaelis-Menten equation was used to obtain the affinity constant and maximum dissociation rate. Various pulling forces can be applied to actin branches as described in (Pandit et al., 2020). The dissociation of actin branches is measured under a range of pulling forces, in the presence or absence of 1 µM VCA or 1 µM GMF (Fig 5A-D). Likewise, the dissociation rate of aged actin branches is performed by exposing branches to 0.15 μM G-actin for 20 minutes prior to monitor their dissociation (Fig 6).
- **Branching efficiency of different NPFs (VCA domain of NWASP WASP, and WASH)**. The coverslip surface of a microfluidics chamber was functionalized by exposing it to 4 pM spectrin-actin seeds for 5 minutes, followed by a passivation with a solution containing 5% BSA for 5 minutes. Mother filaments were polymerized by flowing 0.7 μM 10% Alexa-488 actin. Then for 90 seconds, we exposed the mother filaments to a solution of 20nM Arp2/3, 50nM VCA, and 0.4 μM 10% Alexa-568 actin to generate branches. We next flowed in a solution of 0.5 μM 10% Alexa-568 actin for 90 seconds to further grow the branches. After that time, branch densities were determined by counting the number of branches on mother filaments, and measuring the length of mother filaments.

#### GST pull-down assays

Prior to each pull-down assay, the Glutathione Sepharose 4B resin (GE Healthcare) was incubated with 5% BSA for 5 minutes and washed 3 times with 500 μL GST pull-down buffer containing 50 mM Tris-HCl pH 7.0, 1 mM DTT, 50 mM KCl, and 5% glycerol. 100 μL of 2 μM Glutathione Sepharose 4B attached GST SPIN90 full length was mixed with 250 nM Arp2/3. A concentration gradient (0, 0.5, 2, 4 μM) of GMF or GFP-VCA was mixed with the loaded resin and allow to incubate for 1 hour at 4°C. The resin were then washed 3 times with 300 μL of GST pull-down buffer. Resin-attached proteins were eluted with 50 μL of 20 mM GSH. The sample was separated by SDS-PAGE for western blot analysis. Anti-ArpC2 (Sigma) was used to detect Arp2/3.

The negative control experiment was done in the same way except 50 μL of 2 μM Glutathione Sepharose™ 4B attached GST was mixed with 250 nM Arp2/3, with or without 4 μM GMF or GFP-VCA for 1 hour at 4°C.

### Data analysis

Fiji software (Schindelin et al., 2012) was used to analyze images manually.

#### Survival fraction analysis

We used the Kaplan-Meier method (‘survival’ package in R) to estimate and plot the survival fractions of the different populations probed in microfluidics assays. Data was fitted with a single exponential decay function to obtain the experimental detaching or debranching rates.

In nucleation experiments (Fig. 1), the total number of nucleation sites in the field of view (plateau) was determined by the exponential fit of the curve, and used to normalize the number of nucleated filaments.

#### VCA and GMF protein affinity for SPIN90-Arp2/3 or Arp2/3 branches

The binding affinity curves in figures 2 & 3 were derived using the Michaelis-Menten equation : k_obs(x) = k_max.x/(K_D_+x), where x is the protein concentration in solution, k_max is the maximum apparent rate of the reaction and K_D_ the dissociation constant of the pseudo first-order reaction.

#### Statistical significance

To compare the survival distributions of two samples., p-value from the log-rank test, using the ‘survival’ package in R.

In figure 4, to assess the significance of the statistical difference between fractions of SPIN90-Arp2/3 complexes that renucleated a filament in different conditions (figure 4 B&C), or the fraction of elongating pointed ends after their detachment from SPIN90-Arp2/3 (figure 4 E&F), p-values were calculated based on a chi-square test using R.

## ACKNOWLEDGEMENTS

LC was supported by the European Union’s Horizon 2020 Marie Sklodowka-Curie individual fellowship program (H2020-MSCA-IF-101028239 –MolecularArp). FG was supported by a doctoral grant from the French Ministry of Research. MW was supported by Cancer Research UK (FC001209), the UK Medical Research Council (FC001209), and the Wellcome Trust (FC001209) funding at the Francis Crick Institute as well as by the European Research Council (ERC) under the European Union’s Horizon 2020 research and innovation programme (grant agreement No 810207 to MW). AJ was supported by the ERC (grant StG-679116). GRL was supported by Agence Nationale de la Recherche (grant RedoxActin). For the purpose of Open Access, the authors have applied a CC BY public copyright licence to any Author Accepted Manuscript version arising from this submission.

