## Supplemental figures for "Nucleation and stability of branched versus linear Arp2/3-generated actin filaments"

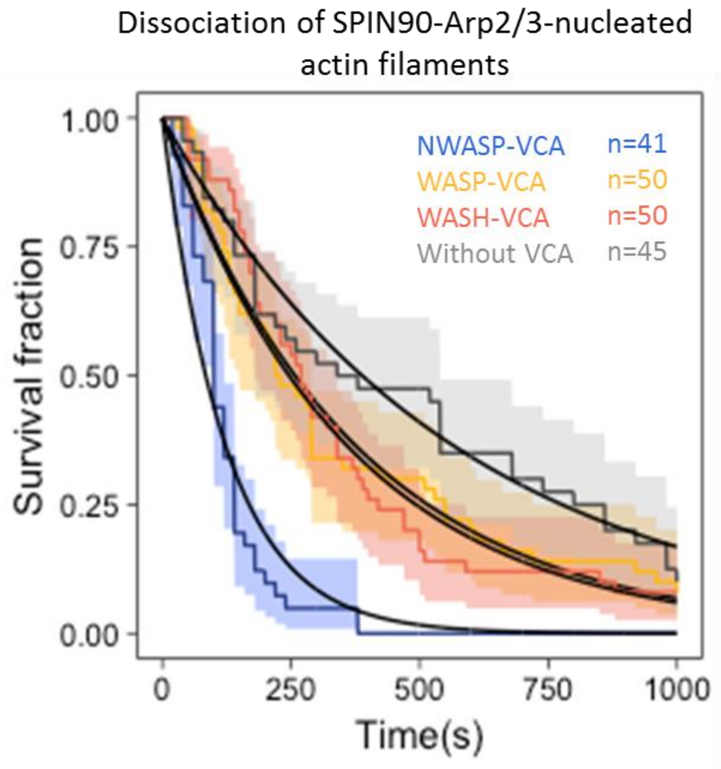


**Figure S1**. Detachment of SPIN90-Arp2/3-nucleated filaments exposed to VCA, during the nucleation experiment shown in figure 1.

Normalized number of filaments dissociated from the surface exposed to 2 μM G-actin (15% labeled with Alexa488) and 1 μM profilin, with 0 or 0.5μM of GST-VCA from different NPFs. Solid lines are exponential fits, yielding dissociation rates $k_{off}$=(1.8±0.6)x$10^{-3}$ $s^{-1}$ without VCA, and $k_{off}$=(7.9±1.9)x$10^{-3}$, (2.7±0.9)x$10^{-3}$, and (2.8±0.7)x$10^{-3}$ $s^{-1}$ with VCA from N-WASP, WASP and WASH, respectively. Indicated values of n are the number of filaments observed in each experiment. These experiments were repeated three times, with similar results.

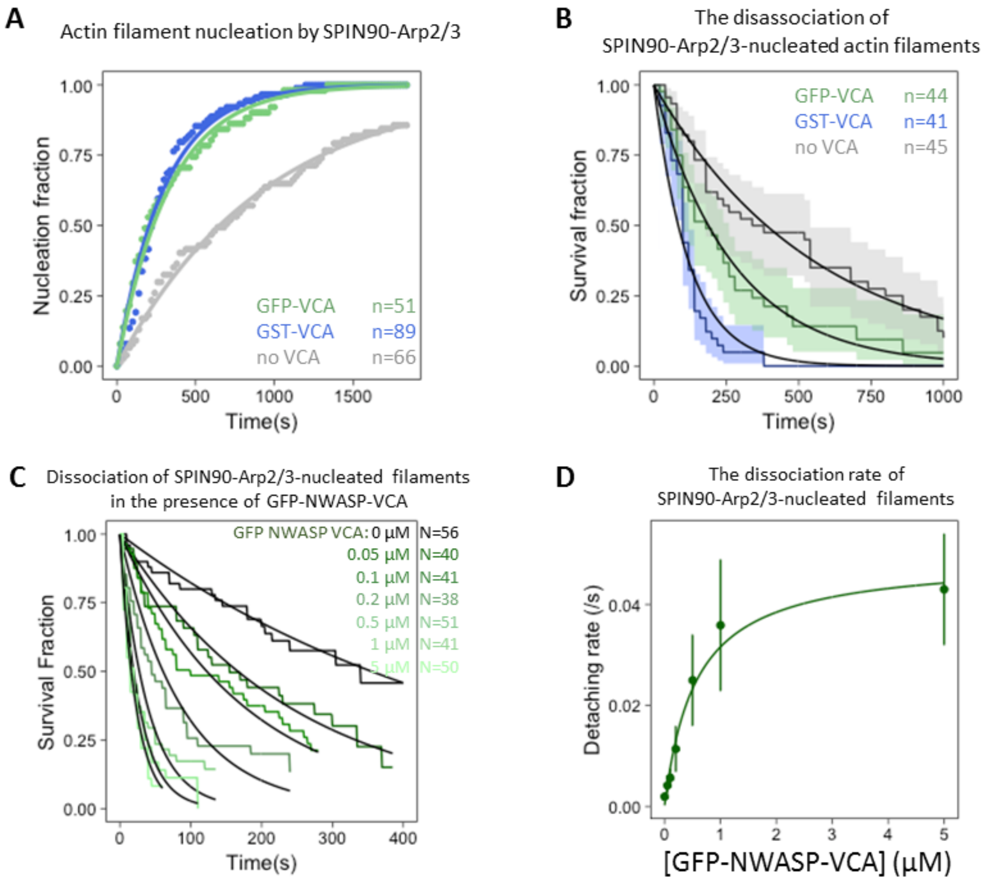


**Figure S2**. Impact of VCA dimerization.

**A.** Normalized number of filaments nucleated over time, from SPIN90-Arp2/3 exposed to 2 µM G-actin (15% labeled with Alexa488) and 1 µM profilin, with 0 or 0.5µM of GST-N-WASP-VCA or GFP-N-WASP-VCA. Solid lines are exponential fits, yielding nucleation rates $k_{nuc}=(1.06\pm0.03)\times10^{-3}\ s^{-1}$ without VCA, and $k_{nuc}=(3.23\pm0.08)\times10^{-3}\ s^{-1}$ and $(3.02\pm0.04)\times10^{-3}\ s^{-1}$ with GST-N-WASP-VCA and GFP-N-WASP-VCA, respectively. Indicated values of n are the number of filaments observed in each experiment. These experiments were repeated three times, with similar results.

**B.** Detachment of SPIN90-Arp2/3-nucleated filaments during the nucleation experiment shown in panel A. Solid lines are exponential fits, yielding dissociation rates $k_{off}=(1.8\pm0.6)\times10^{-3}\ s^{-1}$ without VCA, and $k_{off}=(7.9\pm1.9)\times10^{-3}\ s^{-1}$, $(3.7\pm1.4)\times10^{-3}\ s^{-1}$ with GST-N-WASP-VCA and with GFP-N-WASP-VCA respectively. Indicated values of N are the number of filaments observed in each experiment.

**C.** The fraction of filaments still attached to the surface, versus time, for different concentrations of GFP-NWASP-VCA. Black lines are exponential fits. Indicated values of n are the number of filaments observed in each experiment.

**D.** Detachment rates, determined by exponential fits of survival curves (in C) as a function of the concentration of VCA domains from different NPFs. The data are fitted by a Michaelis-Menten equation, resulting in $K_D$ = 0.56±0.11 µM and $V_{max}$ = 0.049±0.0034 $s^{-1}$.

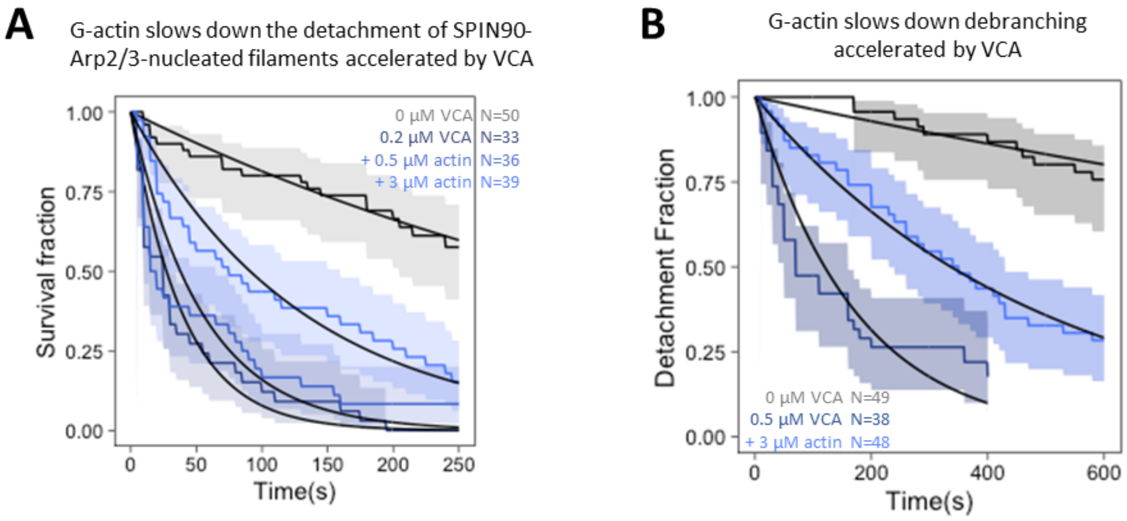


**Figure S3**. The presence of G-actin inhibits the destabilizing action of VCA.

A. The fraction of filaments still attached to the surface versus time, exposed to buffer, or to 0.2 µM VCA with different concentrations of G-actin. Black lines are exponential fits, yielding dissociation rates $k_{off}=(2.1\pm0.8)\times10^{-3}$ $s^{-1}$ without VCA, and $k_{off}=(2.6\pm0.7)\times10^{-2}$ $s^{-1}$ with 0.2 µM VCA, $(1.8\pm0.8)\times10^{-2}$ $s^{-1}$ with 0.2 µM VCA plus 0.5 µM G-actin, $(0.76\pm0.28)\times10^{-2}$ $s^{-1}$ with 0.2 µM VCA plus 3 µM G-actin. The shaded areas represent 95% confidence intervals. Indicated values of n are the number of filaments monitored in each experiment.

B. The fraction of actin branches still attached to the mother filaments versus time, exposed to buffer, or to 0.5 µM VCA without actin, or to 0.5 µM VCA with 3 µM G-actin. Black lines are exponential fits, yielding dissociation rates $k_{off}=(0.37\pm0.7)\times10^{-3}$ $s^{-1}$ without VCA, and $k_{off}=(5.8\pm2.3)\times10^{-3}$ $s^{-1}$ with 0.5 µM VCA, and $(2.1\pm0.7)\times10^{-3}$ $s^{-1}$ with 0.5 µM VCA plus 3 µM G-actin. The shaded areas represent 95% confidence intervals. Indicated values of n are the number of filaments monitored in each experiment. The force applied to the actin branches in this experiment is around 0.2 pN.

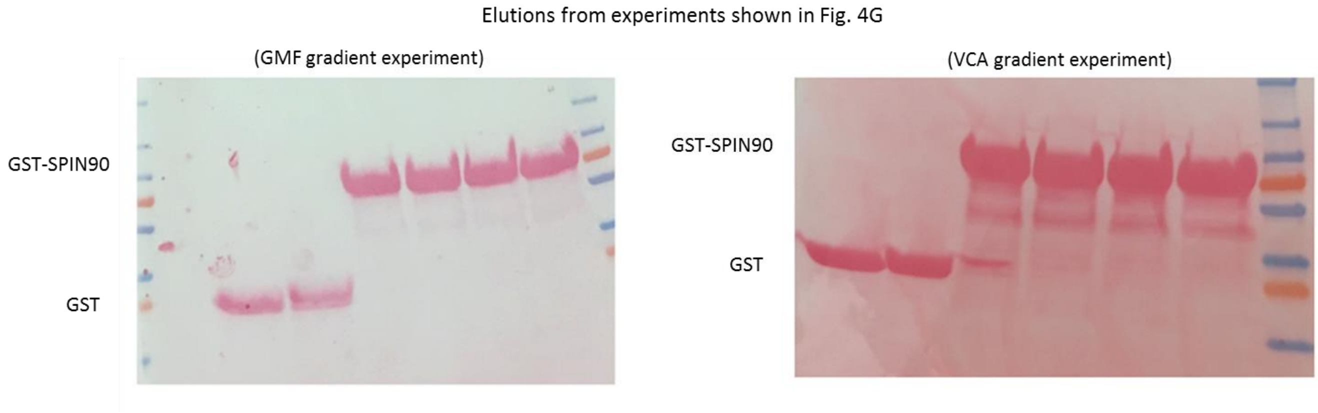


**Figure S4**. Amount of GST and GST-SPIN90 loaded on beads in the pull-down assays (Fig. 4G). In the experiments shown in Fig 4G, before passivating and adding with antibodies, the membrane was stained with Ponceau red.

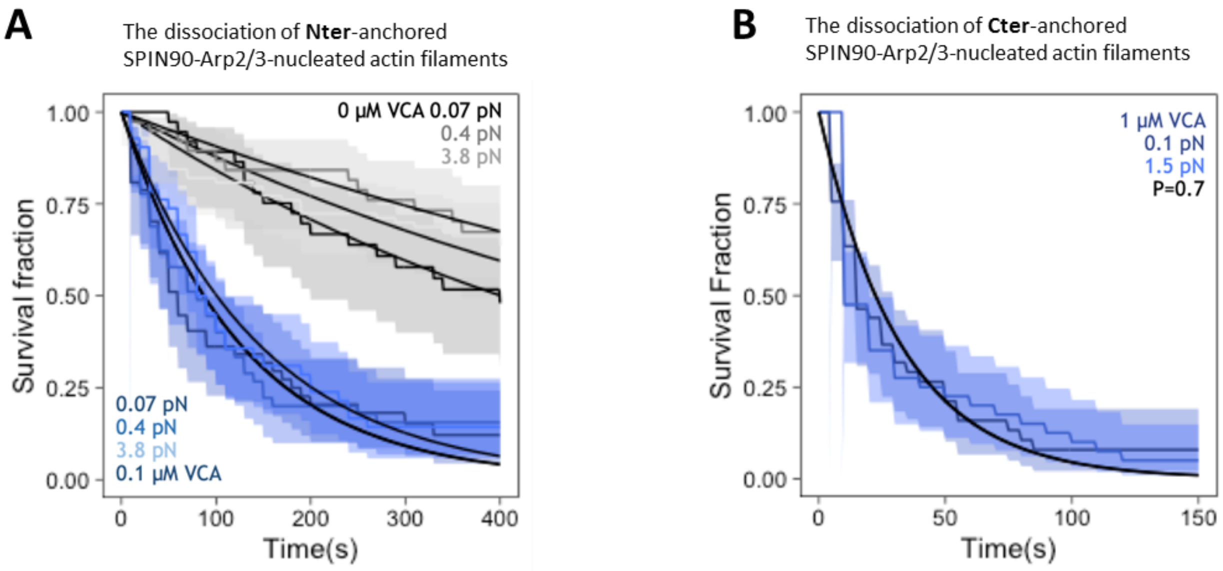


**Figure S5**. Anchoring SPIN90 by its C- or N-terminus has no impact on its behavior.

A. Survival fractions for SPIN90-Arp2/3-nucleated filaments exposed to different forces in the microfluidics chamber, while being exposed to 0.15 µM G-actin alone or with 0.1µM GFP-VCA. In this case, SPIN90 was anchored on the surface through its Nter-GST-tag. Black lines are exponential fits, yielding dissociation rates at 0.07pN force, $k_{off}$=(1.6±0.6)x$10^{-3}$ $s^{-1}$ without VCA, and $k_{off}$=(7.5±3.3)x$10^{-3}$ $s^{-1}$ with 0.1 µM VCA; at 0.4pN force, $k_{off}$=(1.2±0.5)x$10^{-3}$ $s^{-1}$ without VCA, and $k_{off}$=(7.4±3.5)x$10^{-3}$ $s^{-1}$ with 0.1 µM VCA; 3.8pN, $k_{off}$=(1.3±0.4)x$10^{-3}$ $s^{-1}$ without VCA, and $k_{off}$=(6.9±2.4)x$10^{-3}$ $s^{-1}$ with 0.1 µM VCA. The shaded areas represent 95% confidence intervals.

B. Survival fractions for SPIN90-Arp2/3-nucleated filaments exposed to 1 µM VCA. In this case, SPIN90 was anchored on the surface through its Cter-His-tag. Black lines are exponential fits, yielding dissociation rates $k_{off}$=(3.1±1.5)x$10^{-2}$ $s^{-1}$ at 0.1 pN force and $k_{off}$=(3.1±1.3)x$10^{-2}$ $s^{-1}$ at 1.5 pN force. The shaded areas represent 95% confidence intervals.

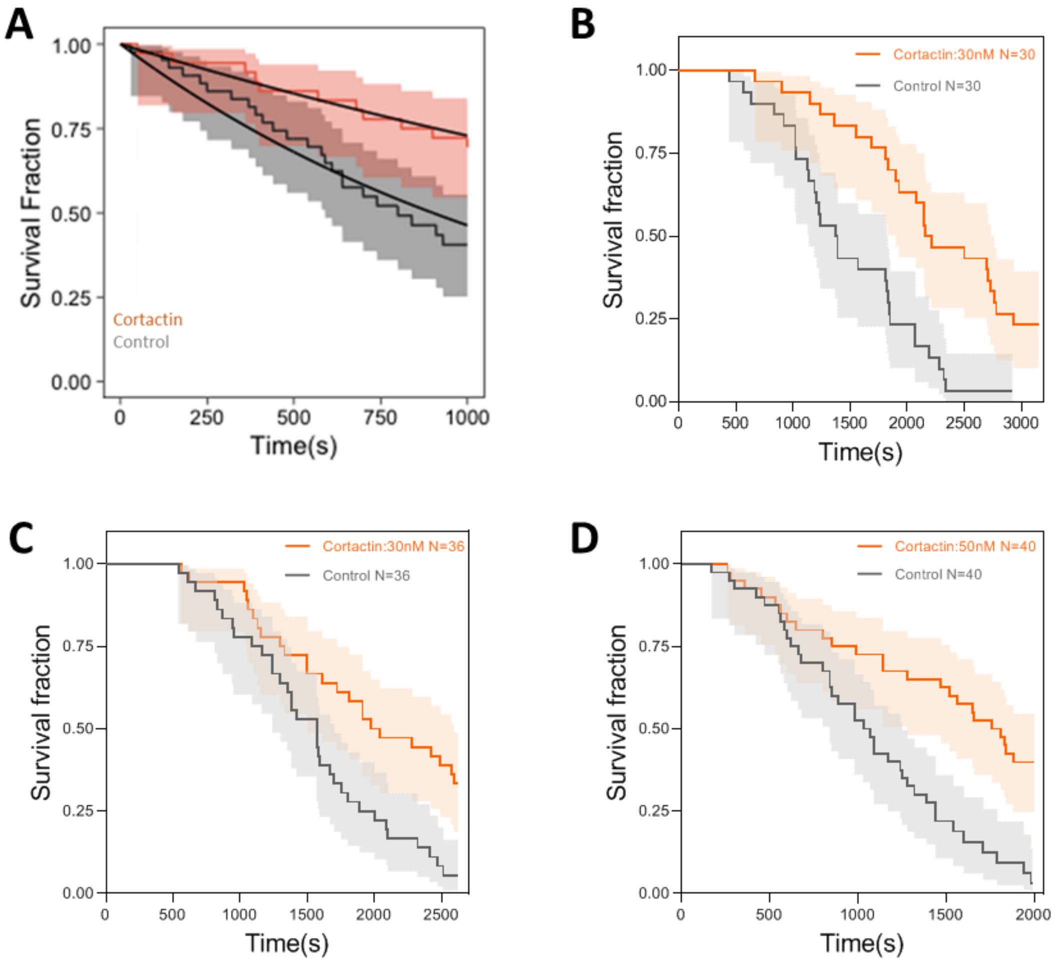


**Figure S6**. Cortactin slows down debranching.

A. The fraction of remaining branches, versus time, exposed to 0.15 µM G-actin (black) supplemented with 100 nM cortactin (red). The branch junctions were exposed to an average force of 0.2 pN. Black lines are exponential fits, yielding dissociation rates at $k_{off}=(7.7\pm1.4)\times10^{-4}\ s^{-1}$ without cortactin, and $k_{off}=(3.2\pm1.5)\times10^{-4}\ s^{-1}$ with 100 nM cortactin.
B. The fraction of remaining branches, versus time, exposed to 0.3 µM G-actin (black) supplemented with 30 nM cortactin (orange). In this case, actin keeps polymerizing during the measurement. Thereby, the forces applied on actin branches increase over time.
C. The fraction of remaining branches, versus time, exposed to 0.18 µM G-actin (black) supplemented with 30 nM cortactin (orange). The branch junctions were exposed to an average force of 0.38 pN, after being aged for 5 minutes in the absence of force.
D. The fraction of remaining branches, versus time, exposed to 0.15 µM G-actin (black) supplemented with 50 nM cortactin (orange). The branch junctions were exposed to an average force of 0.68 pN, after being aged for 10 minutes in the absence of force.

(A-D) The shaded areas represent 95% confidence intervals.

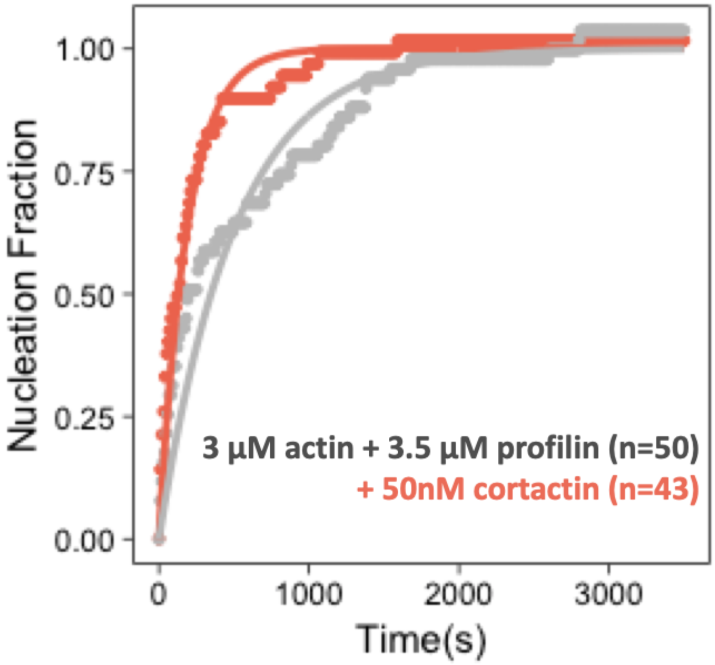


**Figure S7**. SPIN90-Arp2/3 filament nucleation is accelerated by cortactin.
Normalized number of filaments nucleated over time, from SPIN90-Arp2/3 exposed to 3 µM G-actin (15% labeled with Alexa488) and 3.5 µM profilin, with 0 or 50 nM of cortactin. Solid lines are exponential fits, yielding nucleation rates $k_{nuc}$=(2.0±0.04)x$10^{-3}$ $s^{-1}$ without cortactin, and $k_{nuc}$=(5.3±0.08)x$10^{-3}$ $s^{-1}$ with cortactin. Indicated values of n are the number of filaments observed in each experiment..
